# 290 Metagenome-assembled Genomes from the Mediterranean Sea: a resource for marine microbiology

**DOI:** 10.1101/069484

**Authors:** Benjamin J. Tully, Rohan Sachdeva, Elaina D. Graham, John F. Heidelberg

## Abstract

The *Tara Oceans* Expedition has provided large, publicly-accessible microbial metagenomic datasets from a circumnavigation of the globe. Utilizing several size fractions from the samples originating in the Mediterranean Sea, we have used current assembly and binning techniques to reconstruct 290 putative high-quality metagenome-assembled bacterial and archaeal genomes, with an estimated completion of ≥50%, and an additional 2,786 bins, with estimated completion of 0-50%. We have submitted our results, including initial taxonomic and phylogenetic assignments, for the putative high-quality genomes to open-access repositories for the scientific community to use in ongoing research.

## Introduction

Microorganisms are a major constituent of the biology within the world’s oceans and act as the important linchpins in all major global biogeochemical cycles^1^. Marine microbiology is among the disciplines at the forefront of advancements in understanding how microorganisms respond to and impact the local and large-scale environments. An estimated 10^29^ Bacteria and Archaea^2^ reside in the oceans and represent an immense amount of poorly constrained, and ever evolving genetic diversity.

The *Tara Oceans* Expedition (2003-2010) encompassed a major endeavor to add to the body of knowledge collected during previous global ocean surveys to sample the genetic potential of microorganisms^3^. To accomplish this goal, *Tara Oceans* sampled planktonic organisms (viruses to fish larvae) at two major depths, the surface ocean and the mesopelagic. The amount of data collected was expansive and included 35,000 samples from 210 ecosystems^3^. The *Tara Oceans* Expedition generated and publically released 7.2 Tbp of metagenomic data from 243 ocean samples from throughout the global ocean, specifically targeting the smallest members of the ocean biosphere, the viruses, Bacteria and Archaea, and picoeukaryotes^4^. Initial work on these fractions produced a large protein database, totaling >40 million nonredundant protein sequences and identified >35,000 microbial operational taxonomic units (OTUs)^4^.

Leveraging the publically available metagenomic sequences from the “girus” (giant virus; 0.22-1.6 μm), “bacteria” (0.22-1.6 μm), and “protist” (0.8-5 μm) size fractions, we have performed a new joint assembly of these samples using current sequence assemblers (Megahit^5^) and methods (combining assemblies from multiple sites using Minimus2^6^). These metagenomic assemblies were binned using BinSanity^7^ in to 290 high-quality (low contamination) microbial genomes, ranging from 50-100% estimated completion. Environmentally derived genomes are imperative for a number of downstream applications, including comparative genomes, metatranscriptomics, and metaproteomics. This series of genomic data can allow for the recruitment of environmental “-omic” data and provide linkages between functions and phylogenies. This method was initially performed on the seven sites from the Mediterranean Sea containing microbial metagenomic samples (TARA007, -009, -018, -023, -025 and -030), but will continue through the various Longhurst provinces^8^ sampled during the *Tara Oceans* project (Figure 1). All of the assembly data is publically available, including the initial Megahit assemblies for each site from the various size fractions and depths and putative (minimal quality control) genomes.

**Figure 1.**
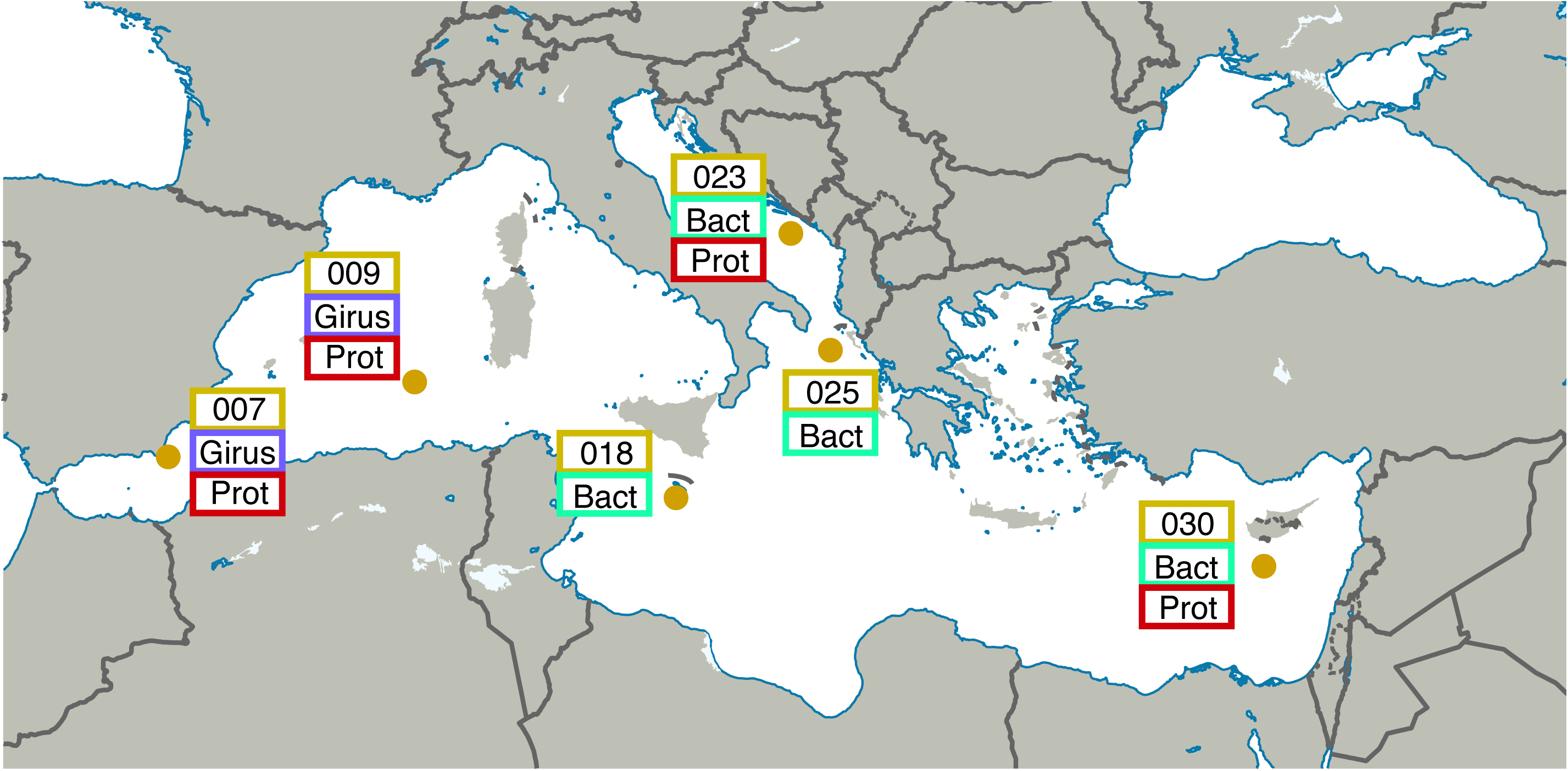
Map illustrating the locations and size fractions sampled for the *Tara Oceans* Mediterranean Sea datasets. Girus, ‘giant virus’ size fraction (0.22-1.6 μm). Bact, ‘bacteria’ size fraction (0.22-1.6 μm). Prot, ‘protist’ size fraction (0.8-5.0 μm).

## Materials and Methods

A generalized version of the following workflow is presented in Figure 2.

**Figure 2.**
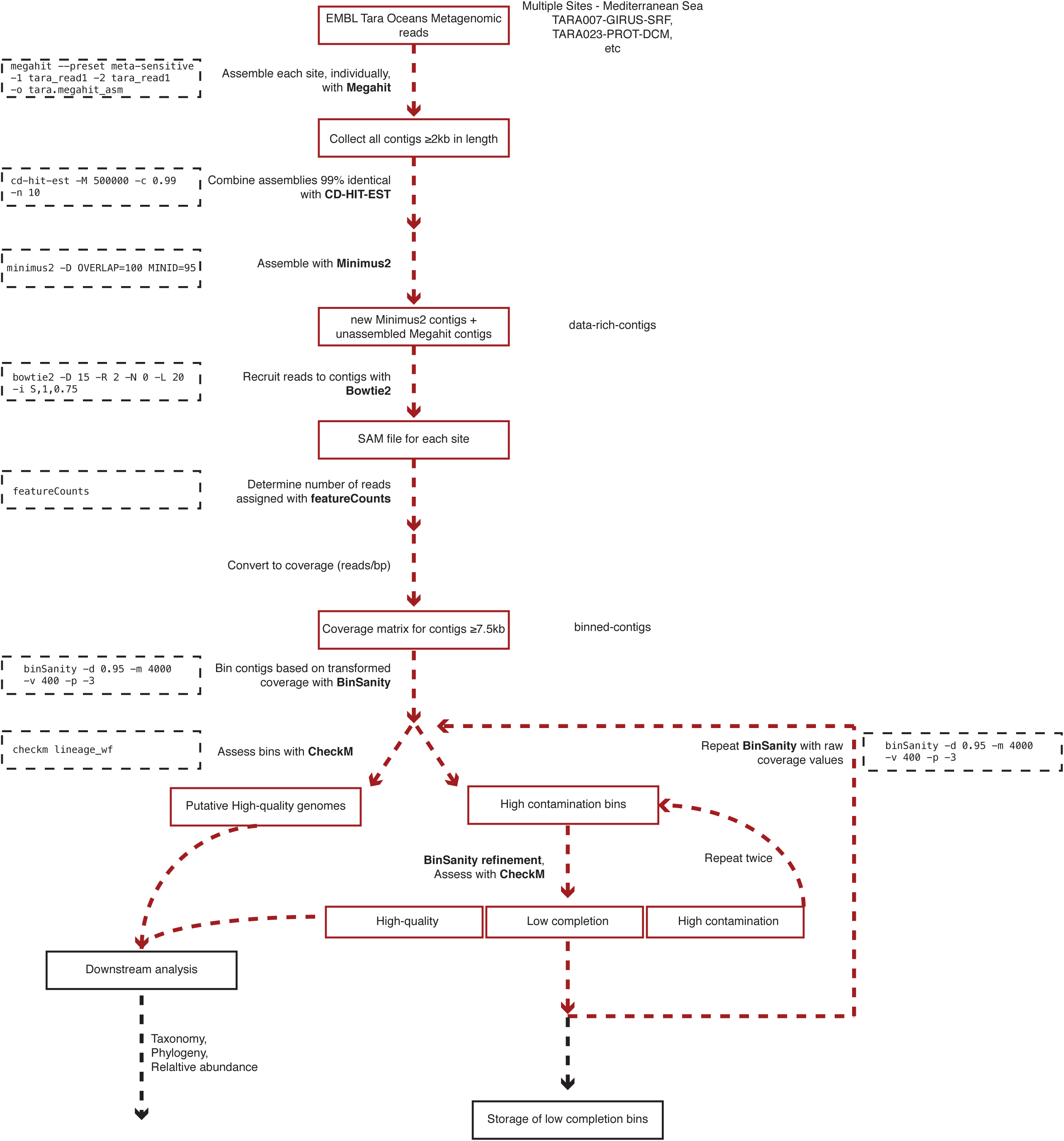
Workflow used to process *Tara Oceans* Mediterranean Sea metagenomic datasets.

### Sequence Retrieval and Assembly

All sequences for the reverse and forward reads from each sampled site and depth within the Mediterranean Sea were accessed from European Molecular Biology Laboratory (EMBL) utilizing their FTP service (Table 1). Paired-end reads from different filter sizes from each site and depth (e.g., TARA0007, girus filter fraction, sampled at the deep chlorophyll maximum) were assembled using Megahit^5^ (v1.0.3; parameters: --preset, meta-sensitive). To keep consistent with TARA sample nomenclature, “bacteria” or “BACT” will be used to encompass the size fraction 0.22-1.6 μm. All of the Megahit assemblies were pooled in to two tranches based on assembly size, ≤1,999bp, and ≥2,000bp. Longer assemblies (≥2kb) with ≥99% semi-global identity were combined using CD-HIT-EST (v4.6; -T 90 -M 500000 -c 0.99 -n 10). The reduced set of contiguous DNA fragments (contigs) was then cross-assembled using Minimus2^6^ (AMOS v3.1.0; parameters: -D OVERLAP=100 MINID=95).

**Table 1.** Statistics for Megahit assemblies, recruitment to data-rich-contigs, and relative abundance of high-quality genome results for each sample

| TARA Sample Site | Size Fraction (Girus, Bacteria, or Protist) | Depth (Surface or DCM*) | No. of reads | No. of initial Megahit assembly | N50<sup>a</sup> (bp; initial Megahit assembly) | Longest initial Megahit assembly (bp) | Recruitment (% data-rich-contigs) | Relative abundance<sup>a</sup> of high-quality genomes (%) | Relative abundance<sup>a</sup> of ten most abundant genomes (%) |
| --- | --- | --- | --- | --- | --- | --- | --- | --- | --- |
| TARA007 | Girus | DCM | 178,519,830 | 1,318,470 | 828 | 220,754 | 72.84 | 14.64 | 6.35 |
| TARA007 | Girus | Surface | 221,166,612 | 1,308,847 | 861 | 211,946 | 81.74 | 14.83 | 6.12 |
| TARA007 | Protist | DCM | 744,458,992 | 4,667,618 | 654 | 188,635 | 19.45 | 8.60 | 3.18 |
| TARA007 | Protist | Surface | 265,432,098 | 2,590,120 | 564 | 18,444 | 25.58 | 1.57 | 0.61 |
| TARA009 | Girus | DCM | 416,553,274 | 2,796,841 | 831 | 1,643,839 | 69.48 | 14.16 | 6.32 |
| TARA009 | Girus | Surface | 489,617,426 | 1,787,467 | 929 | 1,142,851 | 68.85 | 12.29 | 4.76 |
| TARA009 | Protist | DCM | 329,036,110 | 1,938,636 | 613 | 95,724 | 22.07 | 13.35 | 4.20 |
| TARA009 | Protist | Surface | 370,813,078 | 1,700,350 | 588 | 292,050 | 22.53 | 15.97 | 6.17 |
| TARA018 | Bacteria | DCM | 408,021,182 | 2,520,645 | 840 | 1,573,060 | 76.22 | 11.49 | 3.18 |
| TARA018 | Bacteria | Surface | 414,976,308 | 2,604,031 | 816 | 2,086,508 | 75.80 | 11.03 | 3.02 |
| TARA023 | Bacteria | DCM | 147,400,552 | 1,273,576 | 830 | 213,456 | 76.08 | 13.29 | 4.09 |
| TARA023 | Bacteria | Surface | 149,566,010 | 1,237,617 | 825 | 134,179 | 75.98 | 13.82 | 4.01 |
| TARA023 | Protist | DCM | 508,610,652 | 2,707,801 | 734 | 336,689 | 28.23 | 25.07 | 7.83 |
| TARA023 | Protist | Surface | 397,044,232 | 2,246,571 | 593 | 397,140 | 23.00 | 25.16 | 10.31 |
| TARA025 | Bacteria | DCM | 386,627,816 | 2,516,865 | 806 | 388,546 | 69.77 | 14.55 | 5.35 |
| TARA025 | Bacteria | Surface | 457,560,422 | 2,326,838 | 857 | 330,773 | 75.57 | 10.99 | 3.18 |
| TARA030 | Bacteria | DCM | 346,837,034 | 1,968,945 | 1097 | 508,775 | 80.16 | 10.31 | 2.57 |
| TARA030 | Bacteria | Surface | 478,785,582 | 1,639,697 | 1194 | 204,976 | 77.70 | 7.26 | 2.64 |
| TARA030 | Protist | DCM | 426,896,616 | 1,620,343 | 616 | 478,892 | 15.12 | 17.83 | 5.13 |
| TARA030 | Protist | Surface | 430,029,974 | 1,838,588 | 628 | 287,782 | 22.36 | 17.60 | 6.73 |

\*DCM - deep chlorophyll maximum
<sup>a</sup>N50 - length of DNA sequence above which 50% of the total is contained<sup>a</sup>relative abundance - determined using the reads recruited data-rich-contigs

### Metagenome-assembled Genomes

Sequence reads were recruited against a subset of contigs (≥7.5kb) constructed during the secondary assembly (Megahit + Minimus2) for each of the *Tara* samples using Bowtie2^9^ (v4.1.2; default parameters). Utilizing the SAM file output, read counts for each contig were determined using featureCounts^10^ (v1.5.0; default parameters). Coverage was determined for all contigs by dividing the number of recruited reads by the length of the contig (reads/bp). Due to the low coverage nature of the samples, in order to effectively delineate between contig coverage patterns, the coverage values were transformed by multiplying by five (determined through manual tuning). Transformed coverage values were then utilized to cluster contigs in to bins utilizing BinSanity^7^ (parameters: -p -3, -m 4000, -v 400, -d 0.9). Bins were assessed for the presence of putative microbial genomes using CheckM^11^ (v1.0.3; parameters: lineage_wf). Bins were split in to three categories: (1) putative high quality genomes (≥50% complete and ≤10% cumulative redundancy [% contamination – (% contamination × % strain heterogeneity ÷ 100)]); (2) bins with “high” contamination (≥50% complete and ≥10% cumulative redundancy); and (3) low completion bins (<50% complete).

The high contamination bins containing approximately two genomes, three genomes, or ≥4 genomes used the BinSanity refinement method (Binsanity-refine; -m 2000, -v 200, -d 0.9) with variable preference values (-p) of -1000, -500, and -100, respectively. The resulting bins were added to one of the three categories: putative high quality genomes, high contamination bins, and low completion bins. The high contamination bins were processed for a third time with the Binsanity-refine utilizing a preference of -100 (-p -100). These bins were given final assignments to either the putative high quality genomes (some putative genomes had >10% cumulative contamination, but have been designated) or low completion bins.

Any contigs not assigned to putative high-quality genomes were assessed using BinSanity using raw coverage values. Two additional rounds of refinement were performed with the first round of refinement using preference values based on the estimated number of contaminating genomes (as above) and the second round using a set preference of -10 (-p -10). Following this binning phase, contigs were assigned to high quality bins (*e.g.*, ***T****ara* **Med**iterranean genome 1, referred to as TMED1, etc.), low completion bins with at least five contigs (0-50% complete; TMEDlc1, etc. lc, **l**ow **c**ompletion), or were not placed in a bin (Supplemental Table 1 & 2).

**Table 2.** Assembly statistics at various steps during processing

| Contig Grouping | No. of contigs | N50* | Total sequence (bp) |
| --- | --- | --- | --- |
| Megahit assemblies 200-499bp | 24,999,285 | n.d. | 9,293,098,676 |
| Megahit assemblies 500-1,999bp | 16,103,221 | n.d. | 13,382,057,993 |
| Megahit assemblies $\geq 2$kb | 1,517,360 | 4,658 | 6,691,877,664 |
| Megahit assemblies $\geq 2$kb (post-CD-HIT-EST) | 1,126,975 | 4,520 | 4,894,479,496 |
| Minimus2 contigs | 158,414 | 15,394 | 1,727,079,865 |
| Minimus2 + unassembled Megahit contigs $\geq 2$kb (data-rich-contigs) | 660,937 | 5,466 | 3,612,405,904 |
| Minimus2 + unassembled Megahit contigs $\geq 7.5$kb (binned-contigs) | 95,506 | 20,556 | 1,725,063,313 |

\*N50 - length of DNA sequence above which 50% of the total is contained

### Taxonomic and Phylogenetic Assignment of High Quality Genomes

The bins representing the high quality genomes were assessed for taxonomy and phylogeny using multiple methods to provide a quick reference for selecting genomes of interest. Taxonomy as assigned using the putative placement provided via CheckM during the pplacer^12^ step of the analysis to the lowest taxonomic placement (parameters: tree_qa -o 2). This step was also performed for all low completion bins.

Two separate attempts were made to assign the high quality genomes a phylogenetic assignment. High quality genomes were searched for the presence of the full-length 16S rRNA gene sequence using RNAmmer^13^ (v1.2; parameters: -S bac -m ssu). All full-length sequences were aligned to the SILVA SSU reference database (Ref123) using the SINA web portal aligner^14^ (<u>https://www.arb-silva.de/aligner/</u>). These alignments were loaded in to ARB^15^ (v6.0.3), manually assessed, and added to the non-redundant 16S rRNA gene database (SSURef123 NR99) using ARB Parsimony (Quick) tool (parameters: default). A selection of the nearest neighbors to the *Tara* genome sequences were selected and used to construct a 16S rRNA phylogenetic tree. Genome-identified 16S rRNA sequences and SILVA reference sequences were aligned using MUSCLE^16^ (v3.8.31; parameters: -maxiters 8) and processed by the automated trimming program trimAL^17^ (v1.2rev59; parameters: -automated1). Automated trimming results were assessed manually in Geneious^18^ (v6.1.8) and trimmed where necessary (positions with >50% gaps) and re-aligned with MUSCLE (parameters: -maxiters 8). An approximate maximum likelihood (ML) tree with pseudo-bootstrapping was constructed using FastTree^19^ (v2.1.3; parameters: -nt -gtr -gamma; Figure 3).

**Figure 3.**
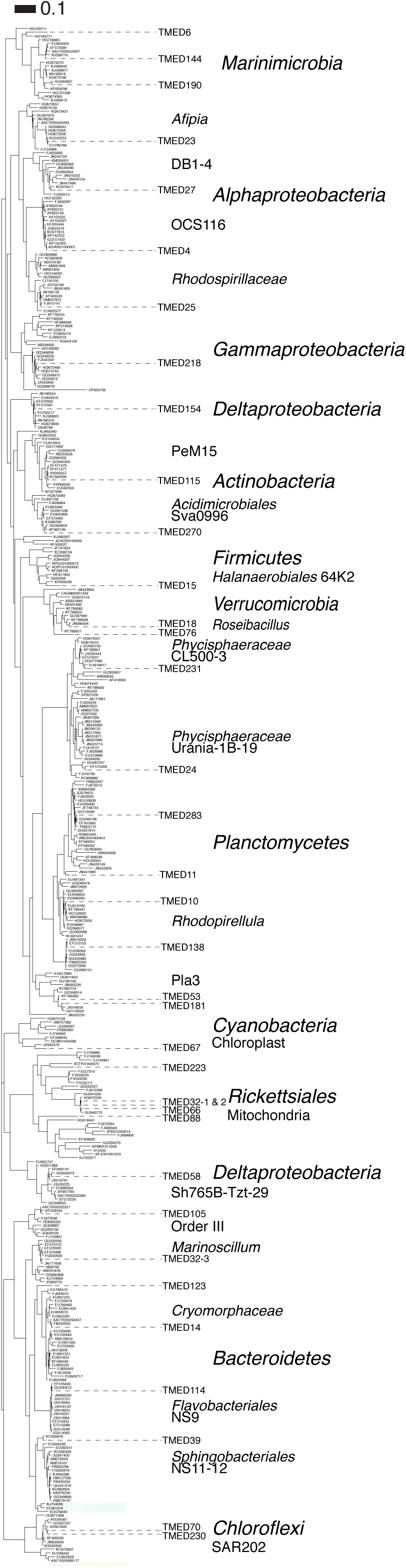
FastTree approximate maximum-likelihood phylogenetic tree constructed with 37 and 785 16S rRNA genes from putative high-quality genomes and references, respectively.

High-quality genomes were assessed for the presence of the 16 ribosomal markers genes used in Hug, *et al*. (2016)^20^. Putative CDSs were determined using Prodigal (v2.6.3; parameters: -m -p meta) and were searched using HMMs for each marker using HMMER^21^ (v3.1b2; parameters: hmmsearch --cut_tc --notextw). If a genome had multiple copies of any single marker gene, neither was considered, and only genomes with ≥8 markers were used to construct a phylogenetic tree. Markers identified from the high quality genomes were combined with markers from 1,729 reference genomes that represent the major bacterial phylogenetic groups (as presented by IMG^22^). Archaeal reference sequences were not included; however, none of the putative archaeal environmental genomes had a sufficient number of markers for inclusion on the tree. Each marker gene was aligned using MUSCLE (parameters: -maxiters 8) and automatically trimmed using trimAL (parameters: -automated1). Automated trimming results were assessed (as above) and re-aligned with MUSCLE, as necessary. Final alignments were concatenated and used to construct an approximate ML tree with pseudo-bootstrapping with FastTree (parameters: -gtr -gamma; Figure 4).

**Figure 4.**
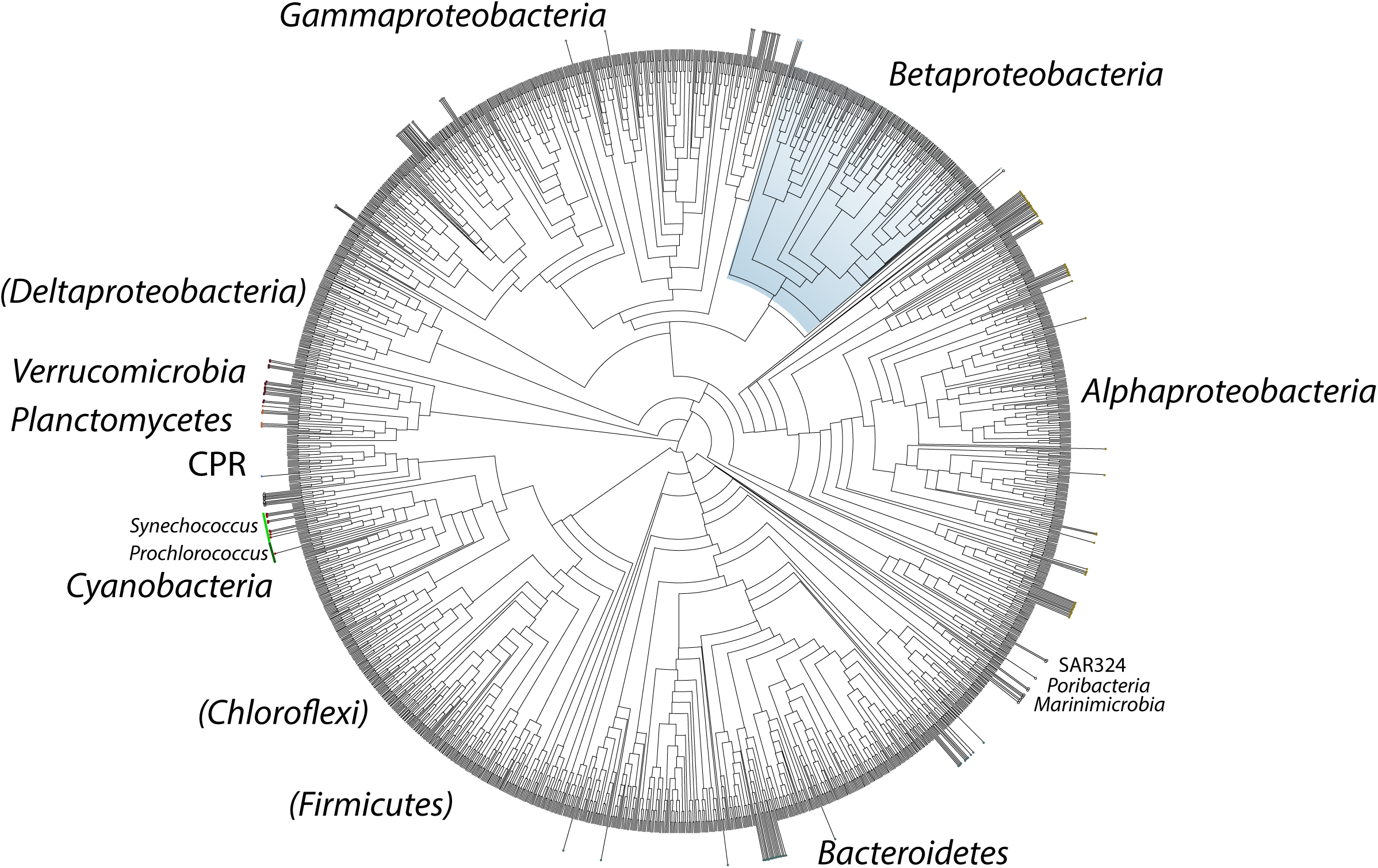
Cladogram of a FastTree approximate maximum-likelihood phylogenetic tree constructed using 16 syntenic, single-copy marker genes for 193 high-quality genomes and 1,729 reference genomes. Leaves denoting the position of the TMED genomes have been indicated by extending beyond the edge of the tree. Sequence alignment is available in Supplemental Information S4. Phylogenetic tree newick file is available in Supplemental Information S5.

### Relative Abundance of High Quality Genomes

To set-up a baseline that could approximate the “microbial” community (Bacteria, Archaea and viruses) present in the various *Tara* metagenomes, which included filter sizes specifically targeting both protists and giruses, reads were recruited against all contigs generated from the Minimus2 and Megahit assemblies ≥2kb using Bowtie2 (default parameters). Some assumptions were made that contigs <2kb would include, low abundance bacteria and archaea, bacteria and archaea with high degrees of repeats/assembly poor regions, fragmented picoeukaryotic genomes, and problematic read sequences (low quality, sequencing artefacts, etc.). All relative abundance measures are relative to the number of reads recruited to the assemblies ≥2kb. Read counts were determined using featureCounts (as above). Length-normalized relative abundance values were determined for each high quality genome for each sample:

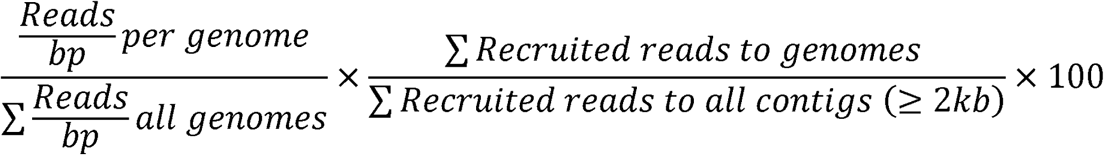

### Data Availability

This project has been deposited at DDBJ/ENA/GenBank under the BioProject accession no. #### and drafts of genomes are available with accession no. #####-#####. Additional files have been provided and are available through FigShare (<u>https://dx.doi.org/10.6084/m9.figshare.3545330</u>), such as: all contigs from Minimus2 + Megahit output used for binning and community assessment; contig read counts per sample; the putative genome contigs and Prodigal-predicted nucleotide and protein putative CDS FASTA files; the ribosomal marker HMM profiles; reference genome markers; high quality genome markers; low completion bins, and contigs without a bin. All contigs generated using Megahit from each sample are available through iMicrobe (http://data.imicrobe.us/project/view/261).

## Results

### Assembly

The initial Megahit assembly was performed on the publicly available reads for *Tara* stations 007, 009, 018, 023, 025, 030. Starting with 147-744 million reads per sample, the Megahit assembly process generated 1.2-4.6 million assemblies with a mean *N*_*50*_ and longest contig of 785bp and 537kb, respectively (Table 1). In general, the assemblies generated from the *Tara* samples targeting the protist size fraction (0.8-5 μm) had a shorter *N*_*50*_ value than the bacteria size fractions (mean: 554bp vs 892bp, respectively). Assemblies from the Megahit assembly process were pooled and separated by length. Of the 42.6 million assemblies generated during the first assembly, 1.5 million were ≥2kb in length (Table 2). Several attempts were made to assemble the shorter contigs, but publicly available overlap-consensus assemblers (Newbler [454 Life Sciences], cap3^23^, and MIRA^24^) failed on multiple attempts. Processing the ≥2kb assemblies from all of the samples through CD-HIT-EST reduced the total to 1.1 million contigs ≥2kb. This group of contigs was subjected to the secondary assembly through Minimus2, generating 158,414 new contigs (all ≥2kb). The secondary contigs were combined with the Megahit contigs that were not assembled by Minimus2. This provided a contig dataset consisting of 660,937 contigs, all ≥2kb in length (Table 2; further referred to as data-rich-contigs).

### Binning

The set of data-rich-contigs was used to recruit the metagenomic reads from each sample using Bowtie2. The data-rich-contigs recruited 15-81% of the reads depending on the sample. In general, the protist size fraction recruited substantially fewer reads than the girus and bacteria size fractions (mean: 19.8% vs 75.0%, respectively) (Table 1). For the protist size fraction, the “missing” data for these recruitments likely results from the poor assembly of more complex and larger eukaryotic genomes. The fraction of the reads that do not recruit in the girus and bacterial size fraction samples could be accounted for by the large number of low quality assemblies (200-500bp) and reads that could not be assembled due to low abundance or high complexity (Table 2).

Unsupervised binning was performed using both transformed and raw coverage values for a subset of 95,506 contigs from the data-rich-contigs that were ≥7.5kb (referred to further as binned-contigs). Binning using the transformed coverage data generated 237 putative high-quality genomes (12 putative genomes are of slightly lower quality with >10% redundancy and have been noted) containing 15,032 contigs (Supplemental Information S1). Contigs not in putative genomes were re-binned based on raw coverage values, generating 53 additional putative high-quality genomes encompassing 3,348 contigs. In total, 290 putative high-quality genomes were generated with 50-100% completion (mean: 69%) with a mean length and number of putative CDS of 1.7Mbp and 1,699, respectively (Supplemental Information S1). All other contigs were grouped in to bins with at least five contigs, but with estimated completion of 0-50% (2,786 low completion bins; 74,358 contigs; Supplemental Information S2) or did not bin (2,732 contigs). Nearly a quarter of the low completion bins (24.7%) have an estimated completion of 0%.

### Taxonomy, Phylogeny, & Potential Organisms of Interest

The 290 putative high-quality genomes had a taxonomy assigned to it via CheckM during the pplacer step. All of the genomes, except for 20, had an assignment to at least the Phylum level, and 83% of the genomes had an assignment to at least the Class level (Supplemental Information S1).

Phylogenetic information was determined for as many genomes as possible. Genomes were assessed for the presence of full-length 16S rRNA genes. In total, 37 16S rRNA genes were detected in 35 genomes. 16S rRNA genes can prove to be problematic during the assembly steps due the high level of conservation that can break contigs^25^ (Figure 3). Additionally, the conserved regions of the 16S rRNA, depending on the situation, can over- or under-recruit reads, resulting in coverage variations that can misplace contigs in to the incorrect genome. As such, several of the 16S rRNA phylogenetic placements support the taxonomic assignments, while some are contradictory. Further analysis should allow for the determination of the most parsimonious result.

Beyond the 16S rRNA gene, genomes were searched for 16 conserved, syntenic ribosomal markers. Sufficient markers (≥8) were identified in 193 of the genomes (67%) and placed on a tree with 1,729 reference sequences (Figure 4). Phylogenies were then assigned to the lowest taxonomic level that could be confidently determined. These putative results reveal a number of genomes were generated that represent multiple clades for which environmental genomic information remains limited, including: *Planctomycetes*, *Verrucomicrobia*, *Marinimicrobia*, *Cyanobacteria*, and uncultured groups within the *Alpha*- and *Gammaproteobacteria*.

### Relative Abundance

A length-normalized relative abundance value was determined for each genome in each sample based on the number of reads recruited to the data-rich-contigs. The relative abundance for the individual genomes was determined based on this portion of the dataset (Supplemental Information S3). In general, the genomes had low relative abundance (maximum relative abundance = 1.9% for TMED155 a putative *Cyanobacteria* at site TARA023 from the protistan size fraction sampled at the surface; Supplemental Table 1). The high-quality genomes accounted for 1.57-25.16% of the approximate microbial community as determined by the data-rich-contigs (mean = 13.69%), with the ten most abundant genomes in a sample representing 0.61-10.31% (Table 1).

## Concluding Statement

The goal of this project was to provided preliminary putative genomes from the *Tara Oceans* microbial metagenomic datasets. The 290 putative high-quality genomes and 2,786 low completion bins were created using the 20 samples and six stations from the Mediterranean Sea.

Initial assessment of the phylogeny of these metagenomic-assembled genomes indicates several new genomes from environmentally relevant organisms, including, approximately 14 new *Cyanobacteria* genomes within the genera *Prochlorococcus* and *Synechococcus* and 33 new SAR11 genomes. Additionally, there are putative genomes from the marine *Euryarchaeota* (n = 11), *Verrucomicrobia* (n = 17), *Planctomycetes* (n = 14), and *Marinimicrobia* (n ≈ 5). Additionally, the low completion bins may house distinct viral genomes. Of particular interest may be the 40 bins with 0% completion (based on single-copy marker genes), but that contain >500kb of genetic material (including 3 bins with >1Mb). These large bins lacking markers may be good candidates for research in to the marine “giant viruses” and episomal DNA sources (plasmids, etc.).

It should be noted, researchers using this dataset should be aware that all of the genomes generated from these samples should be used as a resource with some skepticism towards the results being an absolute. Like all results for metagenome-assembled genomes, these genomes represent a best-guess approximation of a taxon from the environment^26^. Researchers are encouraged to confirm all claims through various genomic analyses and accuracy may require the removal of conflicting sequences.

## Acknowledgements

We are indebted to the *Tara Oceans* project and team for their commitment to open-access data that allows data aficionados to indulge in the data and attempt to add to the body of science contained within. And we thank the Center for Dark Energy Biosphere Investigations (C-DEBI) for providing funding to BJT and JFH (OCE-0939654).

## Author Contributions

BJT conceived of the project, performed all of the methods and analyses, and wrote the manuscript. RS provided the origins of the workflow and invaluable feedback during the execution of the methods and analyses. EDG provided feedback and troubleshooting using the pre-release version of BinSanity. JFH provided funding. RS and JFH contributed to manuscript editing and polishing. All authors have read the submitted draft of the manuscript.

## Supplemental Information

Supplemental Information S1. Statistics, taxonomic and phylogenetic assignments for the putative high-quality genomes.

Supplemental Information S2. Statistics and CheckM taxonomy for low completion bins.

Supplemental Information S3. Relative abundance values determined for each genome based the length-normalized fraction of reads recruited to the genome relative to reads recruited for the data-rich-contigs.

Supplemental Information S4. Concatenated MUSCLE alignment file of 16 ribosomal marker proteins used to construct Figure 4.

Supplemental Information S5. Newick file of concatenated 16 ribosomal marker proteins, including FastTree determined local support values using the Shimodaira-Hasegawa test.

## References

1 Falkowski, P. G., Fenchel, T. & DeLong, E. F. The Microbial Engines That Drive Earth's Biogeochemical Cycles. Science 320, 1034–1039 (2008).

2 Whitman, W. B., Coleman, D. C. & Wiebe, W. J. Prokaryotes: The unseen majority. Proc. Natl. Acad. Sci. U.S.A. 95, 6578–6583 (1998).

3 Karsenti, E. et al. A Holistic Approach to Marine Eco-Systems Biology. Plos Biol 9, e1001177–5 (2011).

4 Sunagawa, S. et al. Ocean plankton. Structure and function of the global ocean microbiome. Science 348, 1261359–1261359 (2015).

5 Li, D. et al. MEGAHIT v1.0: A fast and scalable metagenome assembler driven by advanced methodologies and community practices. Methods 102, 3–11 (2016).

6 Treangen, T. J., Sommer, D. D., Angly, F. E., Koren, S. & Pop, M. Next generation sequence assembly with AMOS. Curr Protoc Bioinformatics Chapter 11, Unit 11.8 (2011).

7 Graham, E. D., Heidelberg, J. F. & Tully, B. J. BinSanity: unsupervised clustering of environmental microbial assemblies using coverage and affinity propagation. PeerJ 5, e3035–19 (2017).

8 Longhurst, A., Sathyendranath, S., Platt, T. & Caverhill, C. An Estimate of Global Primary Production in the Ocean From Satellite Radiometer Data. Journal of Plankton Research 17, 1245–1271 (1995).

9 Langmead, B. & Salzberg, S. L. Fast gapped-read alignment with Bowtie 2. Nat Meth 9, 357–359 (2012).

10 Liao, Y., Smyth, G. K. & Shi, W. featureCounts: an efficient general purpose program for assigning sequence reads to genomic features. Bioinformatics 30, 923–930 (2014).

11 Parks, D. H., Imelfort, M., Skennerton, C. T., Hugenholtz, P. & Tyson, G. W. CheckM: assessing the quality of microbial genomes recovered from isolates, single cells, and metagenomes. Genome Res. 25, 1043–1055 (2015).

12 Matsen, F. A., Kodner, R. B. & Armbrust, E. V. pplacer: linear time maximum-likelihood and Bayesian phylogenetic placement of sequences onto a fixed reference tree. BMC Bioinformatics 11, 538 (2010).

13 Lagesen, K. et al. RNAmmer: consistent and rapid annotation of ribosomal RNA genes. Nucleic Acids Res. 35, 3100–3108 (2007).

14 Pruesse, E., Peplies, J. & Glöckner, F. O. SINA: accurate high-throughput multiple sequence alignment of ribosomal RNA genes. Bioinformatics 28, 1823–1829 (2012).

15 Ludwig, W. et al. ARB: a software environment for sequence data. Nucleic Acids Res. 32, 1363–1371 (2004).

16 Edgar, R. C. MUSCLE: multiple sequence alignment with high accuracy and high throughput. Nucleic Acids Res. 32, 1792–1797 (2004).

17 Capella-Gutiérrez, S., Silla-Martínez, J. M. & Gabaldón, T. trimAl: a tool for automated alignment trimming in large-scale phylogenetic analyses. Bioinformatics 25, 1972–1973 (2009).

18 Kearse, M. et al. Geneious Basic: an integrated and extendable desktop software platform for the organization and analysis of sequence data. Bioinformatics 28, 1647–1649 (2012).

19 Price, M. N., Dehal, P. S. & Arkin, A. P. FastTree 2--approximately maximum-likelihood trees for large alignments. PLoS ONE 5, e9490 (2010).

20 Hug, L. A. et al. A new view of the tree of life. Nature Microbiology 1–6 (2016). doi:10.1038/nmicrobiol.2016.4.

21 Finn, R. D., Clements, J. & Eddy, S. R. HMMER web server: interactive sequence similarity searching. Nucleic Acids Res. 39, W29–W37 (2011).

22 Markowitz, V. M. et al. The integrated microbial genomes (IMG) system. Nucleic Acids Res. 34, D344–8 (2006).

23 Huang, X. & Madan, A. CAP3: A DNA sequence assembly program. Genome Res. 9, 868–877 (1999).

24 Chevreux, B. et al. Using the miraEST assembler for reliable and automated mRNA transcript assembly and SNP detection in sequenced ESTs. Genome Res. 14, 1147–1159 (2004).

25 Miller, J. R., Koren, S. & Sutton, G. Assembly algorithms for next-generation sequencing data. Genomics 95, 315–327 (2010).

26 Sharon, I. & Banfield, J. F. Microbiology. Genomes from metagenomics. Science 342, 1057–1058 (2013).

